# An Empirical Study on Quantifying Physical Energy Expenditure during High-intensity Exercise

**DOI:** 10.1101/2025.10.05.680575

**Authors:** Yu Xing, Xiaojun Zhou, Jian Wang, Jun Zhang, Zhaohui Wang, Hao Yu

## Abstract

In the process of high-intensity exercise (HIE), the accurate evaluation of Physical Energy Expenditure (PEE) is of great significance to many occupational fields. However, due to the limitation of current technology, the quantitative ability of existing PEE evaluation methods in HIE situations is still insufficient. Based on the existing physiological coupling mechanism between cardiovascular parameters and energy metabolism, this study analyzed the dynamic changes of high-frequency blood pressure (BP) and heart rate (HR) during incremental load exercise and high-intensity interval training. The results show that although HR has a nonlinear response to the increase of exercise intensity, its time pattern-when analyzed synergistically with BP fluctuation-can provide reliable metabolic indicators. We used Doubly Labeled Water (DLW) as the gold standard for PEE determination and measured the total energy expenditure (TEE) in three consecutive days. During the monitoring period, high frequency invasive BP and HR monitoring was performed simultaneously, and the time integral value of SBP × HR and its ratio to TEE (i.e. Quotient value) were calculated. By dividing the corresponding SBP × HR integral value during HIE by the above quotient value, the PEE of this exercise can be estimated. Specifically, TEE can be accurately estimated by time integration of the product of SBP and HR. The short-term PEE rate can be solved by the differential equation constructed by the systolic blood pressure(SBP) trend function in this period, and can also be obtained by dividing the PEE in this period by the duration of this period. In this study, based on the results of invasive measurements, a new method for real-time monitoring of HIE using non-invasive cardiovascular parameters during PEE was proposed, providing new ideas and methodological support for applications in related fields.

## 1 Introduction

High-Intensity Exercise (HIE) is usually defined as physical activity with absolute intensity ≥ 6.0 METs, relative intensity ≥ 80% VO_2_max or 77-95% HRmax and subjective feeling RPE15-17[1-4]. It is a strong physiological stimulation factor, which has wide and important significance in competitive sports, physical labor and public health[5-7]. Its remarkable benefit stems from the fact that this exercise mode can cause strong acute metabolic disturbance and long-term chronic adaptive changes. These reactions are different from moderate-intensity continuous training in nature and degree, and often show better promotion effect on various health and physical fitness indicators [8, 9]. Although the amount and rate of PEE during HIE are the key to promote training effect and avoid sports risks, it is always difficult to quantify PEE during HIE due to the short-term nature of HIE and the limitations of existing methods.

### 1.1 Research needs of high-intensity exercise

#### 1.1.1 Competitive sports

HIE has become the core component of competitive athletes’ training plan because of its remarkable effectiveness in rapidly improving anaerobic and aerobic capacity. Research shows that about 89% of competitive sports training programs, especially those that need to repeatedly carry out maximum or near maximum intensity output (such as various team sports, cycling, middle and long distance running, etc.), take HIE as the key training content [10]. The adaptive response induced by this kind of exercise is not limited to the enhancement of metabolic function, but also covers important neuromuscular adaptation, including more efficient recruitment mode of exercise units, stronger explosive force, and good coordination under fatigue [11-13]. These comprehensive improvements in physiological and neuromuscular levels have strengthened the ability of musculoskeletal system to cope with the needs of high-intensity competition, thus significantly reducing the risk of sports injuries [14, 15].

#### 1.1.2 Occupations with high PEE

In some high-risk occupations, developing excellent physical fitness through high-intensity exercise is the core requirement of professional ability. For example, mining operations often need to bear intermittent and extremely strong physical load, and its average task metabolic equivalent (MET) can reach 8.5, which poses a severe challenge to workers’ muscle strength and anaerobic endurance [16, 17]; Forestry workers need to maintain high heart load for a long time at work. Research records show that they can stay at more than 85% of the maximum HR for 72 ± 15 minutes every day. This continuous high-intensity cardiovascular stress state is highly similar to the physiological characteristics of interval training [18, 19]; The training program of special forces is explicitly designed to push the trainees to the physiological limit, which usually includes completing 6 to 8 groups of HIE with an intensity of 90%-100% of the peak oxygen uptake in each class, so as to comprehensively build physical resilience, psychological tolerance and the ability to maintain performance under extreme pressure [20, 21].

#### 1.1.3 Metabolic and physiological adaptation

The main physiological effect of HIE lies in its unparalleled efficiency in stimulating mitochondrial production. Studies have shown that compared with moderate-intensity continuous training with equal exercise load, mitochondrial enzyme activity and density increase by 3.2 times after high-intensity exercise, which is very important for enhancing oxidative capacity and endurance performance [22-24]. At the same time, excessive oxygen consumption increased significantly after exercise, reaching 38 ± 12%. This continuous state of increased metabolic rate promoted a large amount of extra calorie consumption, which was mainly realized by supplementing phosphate reserves, eliminating lactic acid and increasing catecholamine-induced thermogenesis, thus contributing about 450 ± 120 kcal of daily PEE increment [25-27]. However, excessive oxygen consumption after exercise is an index after exercise, which lacks real-time significance for risks during exercise.

#### 1.1.4 Advantages of promoting health

From a clinical perspective, HIE is an efficient non-drug intervention strategy, which can significantly improve metabolic health. A pioneering study found that 12 weeks of HIE intervention can increase insulin sensitivity by 28%, which is mainly attributed to the enhancement of GLUT4 transport capacity in skeletal muscle and the improvement of pancreatic β cell function [28-30]. This mechanism makes HIE a powerful means to prevent and manage type 2 diabetes [31]. In addition, the exercise load and metabolic stress caused by HIE can effectively stimulate the process of bone remodeling, improve bone mineral density [32], and promote the overall improvement of cardiovascular health by enhancing stroke volume and improving vascular endothelial function [33, 34].

#### 1.1.5 Risks of high-intensity exercise

Although HIE is significantly effective in enhancing physical fitness, it also comes with considerable physiological risks that need serious attention. Such activity may lead to poor heart adaptation, including myocardial fibrosis and arrhythmia, and even pathological remodeling of the heart-especially in individuals who have not been adequately screened for health [35, 36]. The musculoskeletal system is also vulnerable to acute injuries, such as muscle strain and ligament rupture, as well as long-term conditions caused by overuse, including stress fractures and degenerative joint diseases [37]. In addition, strenuous exercise can induce significant oxidative stress and temporary immunosuppression, thus increasing susceptibility to infection [38, 39]. Exercise rhabdomyolysis is a particularly serious and potentially life-threatening complication [40]. Therefore, HIE must be carried out according to scientific principles, emphasizing gradual progress, full recovery and professional supervision to reduce these related risks.

The acute risk associated with HIE mainly stems from its extreme requirements for various physiological systems of the body. This intense stress may break through the physiological compensation limit, thus causing a series of short-term health damage [41]. Cardiovascular system is particularly fragile in this process. A large number of clinical and epidemiological evidences show that HIE may significantly increase the risk of acute cardiovascular events (such as myocardial infarction and arrhythmia) [42, 43], and is also an important inducement of acute musculoskeletal injury (including muscle tear, ligament sprain and fracture) [44, 45].

To sum up, the benefits and risks of HIE coexist, and with the increase of exercise intensity, the benefit-risk ratio of exercise also changes. In order to find an acceptable balance point, it is particularly important to quantify physical energy consumption reasonably and accurately. Therefore, PEE and its change rate based on HR and BP, as key parameters reflecting exercise efficiency and safety, deserve further study.

### 1.2 Related Applied Research

Doubly labeled water (DLW) technique is recognized as the gold standard for PEE evaluation because of its excellent validity in energy metabolism research [46-48]. Based on the difference elimination kinetics of ^2^H and ^18^O isotopes, this method achieves 97-99% accuracy in long-term longitudinal monitoring, and can continuously evaluate TEE in 3 to 14 days, making it especially suitable for tests that are inconvenient to impose constraints. Some key achievements have been made in the research of using DLW. Pontzer, et al. proved through cross-population analysis that fat-free body weight predicted TEE through power function (β = 0.83), and further revealed the limited energy distribution relationship between basal metabolic rate (BMR) and active energy expenditure (AEE), which is called limited total energy model [49, 50]. However, Trabulsi reported that changes in dietary major nutrient composition and energy intake may lead to systematic deviation of DLW estimates of up to 12%, especially under high-fat diet conditions, and the ^2^H turnover kinetics changes-this is a key methodological limitation that must be considered in application [51-53].

DLW also plays a central role in the study of body composition-metabolism. Using large-scale anthropometric data, Yao, et al. developed a multiple regression model with FFM as the core predictor (R2 = 0.91), which significantly improved the prediction accuracy of TEE [54]. Yosuke, et al. used DLW as a standard method to correct the systematic error of bioelectrical impedance analysis (BIA), which reduced the deviation of body fat estimation Δ = 3.2 kg, and emphasized its reference value in body composition research [55].

In order to solve the limitations of DLW’s long cycle and applicability, supplementary technologies have been developed. ^13^C-sodium bicarbonate labeling method was in good agreement with indirect calorimetry (R2=0. 964, p < 0.001) [56-58]. Because of its minimally invasive and low requirement for metabolic homeostasis, it is widely used in neonatal and infant metabolism research, with an average measurement error of less than 5%. Although indirect calorimetry is limited to laboratory environment, its high time resolution makes it indispensable in the study of sports energy consumption. Ekelund, U., et al. verified its strong consistency with DLW under steady-state conditions (ICC = 0.89) [59]. However, during high-intensity intermittent exercise, the error rate rose to 8-12% due to the nonlinear increase of lung ventilation and CO_2_ production [60-62].

In a word, the current PEE evaluation methods have their own unique advantages and applicable scope. Although DLW is still irreplaceable in longitudinal monitoring without constraints, future research should focus on integrating multi-technology complementary strategies, combining dynamic modeling and machine learning methods [63-66], to further improve the measurement accuracy and applicability across diverse scenes.

### 1.3 Technical bottleneck of detecting PEE during HIE

Under the condition of HIE, the accurate evaluation of PEE faces many technical bottlenecks. First of all, the DLW test period is long, generally 3-14 days, and the average daily PEE is calculated. Although the test conditions are not limited, the PEE of HIE in a short time cannot be accurately measured [67]. Secondly, the measurement error of respiratory quotient (RQ) increased significantly, mainly due to the interference of carbon dioxide retention, unsteady ventilation dynamics and anaerobic glycolysis metabolic pathway [68, 69]. Thirdly, the performance of the energy prediction model based on heart rate decreased significantly, and its correlation coefficient with actual PEE decreased to r < 0.6, reflecting the decoupling between cardiovascular response and real metabolic demand under extreme load, which may be related to the compensatory regulation mechanism triggered by autonomic nervous system under extreme load [70,71].

In actual combat environments such as competitive sports and military training, real-time energy monitoring usually requires the system to have sub-second response capability. However, the existing mainstream wearable devices are limited by the low sampling frequency (usually only 0.5-1 Hz), which makes it difficult to accurately capture the rapid metabolic transformation process in intermittent HIE [72, 73].

In addition, the physiological variability between individuals further increases the complexity of monitoring. Among athletes, the prediction deviation of commonly used algorithms can reach 10-15%, which is partly due to adaptive changes such as specific regulation of autonomic nervous system and remodeling of heart structure caused by long-term training [74, 75]. Environmental factors can also significantly affect measurement accuracy. For example, an increase of 5 °C in ambient temperature may lead to blood flow redistribution caused by thermoregulation, resulting in an additional 3-5% error in energy expenditure estimation [76, 77].

These limitations, which span physiological, technical and environmental fields, together restrict the ecological validity and practical value of the current PEE monitoring system in HIE.

It is worth noting that the dynamic change trend of interventional SBP in HR detection is highly correlated with the PEE of the body, which provides a new breakthrough direction for non-invasive monitoring of exercise load. Further research shows that the integrated analysis of multidimensional cardiovascular parameters (such as stroke volume, arterial pressure waveform characteristics, etc.) can significantly improve the accuracy of sports energy expenditure and load evaluation, thus laying a solid theoretical foundation for building a new generation of athletes’ physical energy evaluation system based on multimodal physiological information.

## 2 Research hypotheses

### 2.1 Dynamic response ability of cardiovascular system

The cardiovascular monitoring framework proposed in the above studies is based on two basic physiological mechanisms: hemodynamic coupling and metabolic feedback regulation [78, 79]. From the hemodynamic point of view, HIE causes a significant increase in cardiac output, accompanied by a significant decrease in systemic peripheral vascular resistance [80]. This regulation mechanism originates from vasodilation of skeletal muscle vascular bed and compensatory vasoconstriction of visceral circulation [81, 82]. This blood flow redistribution significantly improves the efficiency of oxygen delivery, and directly reflects the dynamic response ability of cardiovascular system to metabolic demand [83, 84].

### 2.2 Metabolic rate and cardiovascular capacity

At the metabolic level, a quantitative model of hormone-vessel interaction was established by using the strong correlation between catecholamine release (especially norepinephrine) and systolic blood pressure fluctuation (r = 0.73)[85, 86]. In addition, by accurately capturing the time synchronization between ventilation thresholds (VT1 and VT2) and cardiovascular parameters (such as HR variability and BP inflection point), the recognition ability of metabolic-cardiovascular coupling is significantly enhanced [87-89]. This integrated multi-system physiological modeling method effectively overcomes the limitations of traditional single-parameter monitoring under high physiological load, thus providing a solid physiological foundation for accurately estimating PEE during HIE [90-92].

### 2.3 Physiological mechanism of detecting PEE based on BP + HR

The fundamental principle of HR monitoring for estimating EE is the intrinsic relationship between cardiovascular response and metabolic demand. As the core pump for delivering oxygen and metabolic substrates to tissues throughout the body, cardiac function can indirectly reflect overall energy utilization. Cardiac function indicators used to assess the heart’s pumping capacity, such as ejection fraction (EF), cardiac output (CO), stroke volume (SV), and cardiac index (CI), present significant challenges to measure during human physical activity. Similarly, BP, which reflects the intensity of cardiac work along with HR, is an indicator formed by the combined effect of cardiac contractility and peripheral resistance. The coordinated interaction between HR and BP ensures the stability of blood supply.

Hypothesis: Under normal conditions of cardiac structure, electrical activity, and blood hormones, the levels of BP and HR jointly reflect the metabolic status of the body.

## 3 Method

### 3.1 Participant criteria

A total of 39 participants (31 males and 8 females) were recruited in this study. Among them, 32 students majoring in physical education (average age: 19 ± 0.5 years old, height: 182 ± 5 cm, including 24 national second-class athletes), and 7 members of provincial long jump teams (average age: 19 ± 0.8 years old, height: 184 ± 7 cm, including 3 national first-class athletes and 1 national athlete).

Exclusion: Can’t take part in high-intensity exercise.

### 3.2 Testing equipment

On the athletics field, HR during exercise was recorded using Polar Team II, which has an effective radius of at least 150 meters, a maximum sampling frequency of 1 Hz, and can accommodate up to 28 participants. In the physical training room, two BP testing devices were used: (1) Suntech Tango M2, which can set the collection frequency and complete a measurement every 30 seconds when necessary; (2) Mindray BeneVision N15, which measures BP and HR with a minimum sampling rate of 100 Hz. The N15 is an invasive BP monitoring device that can simultaneously track systolic pressure (SBP), diastolic pressure (DBP), heart rate (HR), tissue oxygenation, and provides an average arterial pressure tracking curve.

### 3.3 Test content

#### 3.3.1 Power bicycle constant load test

In the power bicycle test, the load is set to 100 W and 150 W respectively. Participants pedaled at 40, 60, 80 and 100 rpm for 60 seconds and 120 seconds respectively, and the interval between the two tests was based on the HR returning to the baseline level. 24 participants took part in this project,in this project, the participants were recorded for HR and non-invasive BP data.

#### 3.3.2 Power bicycle maximum load test

Eight national second-class athletes (male) participated in the maximum load test on a power bicycle, during which invasive BP and HR were measured. After wearing the equipment, the participants waited on the power bicycle for a period of time until their HR and BP stabilized, then began pedaling at maximum intensity until they could no longer continue. Subsequently, they rested until reaching a quiet state, and repeated high-intensity pedaling until their BP during the recovery period dropped to a certain value.

#### 3.3.3 Test arrangements for physical education majors

A total of 20 participants (including 4 females) took part in the track tests, all of whom were national second-class athletes. A total of 32 running tests were completed: 60-meter sprints in the first week and 100-meter sprints in the second week. In this project, only HR data of the participants were recorded.

#### 3.3.4 Long jump athletes test schedule

The field running test includes 8 sprints of 60 meters and 100 meters, which are performed every two weeks, and full rest is guaranteed after each test. One of the seven-member teams (including one national athlete and three national first-class athletes) completed eight repeated runs at each distance. In this project, only HR data of the participants were recorded.

### 3.4 Testing conditions and data analysis methods

Data collection occurred during participants’ physical training sessions without disrupting their regular class procedures. Participants understood and approved the content and significance of the data collection and received appropriate compensation.

HR was continuously monitored throughout the whole process, and invasive BP was measured in 8 subsets of participants. The TEE of individuals in three consecutive days was measured by DLW technique. During this period, non-invasive BP and HR were monitored simultaneously, and the integral of the product of systolic blood pressure and heart rate (SBP × HR) with time was calculated. The ratios (denoted as R1, R2, R3) of the integral to the daily energy expenditure (EE) obtained by DLW were calculated respectively, and the degree of dispersion was evaluated. The ratio (R) of the three-day total integral to the TEE obtained by DLW is further determined. Based on the ratio R, the TEE of any time period can be estimated by dividing the SBP × HR integral value by R; Divided by the duration, the average PEE rate in this period can be obtained. In addition, the differential equation constructed by the trend function of SBP in this period can be used to obtain a more accurate instantaneous PEE rate.

## 4 Results

During pedaling exercise, SBP showed significant exercise-induced fluctuation, and its amplitude reached 140% of the baseline level before exercise. As shown in Fig. 1, BP rapidly drops to the lowest point after exercise, showing a multiphase recovery mode: SBP decreases by 5-10% in turn, and diastolic blood pressure (DBP) decreases by about 5% on average, indicating that there is a complex cardiovascular compensatory mechanism after HIE. It is worth noting that HR dynamics show different patterns: there is no statistically significant time trend (p > 0.05) during the whole observation period, and its variability always remains within the typical resting fluctuation range. This separation between BP recovery and cardiac chronotropic response implies the different participation of different branches of autonomic nervous system in the recovery process, and highlights the possible decoupling phenomenon between sympathetic-led vascular regulation and parasympathetic-mediated cardiac reactivation.

**Fig. 1.**
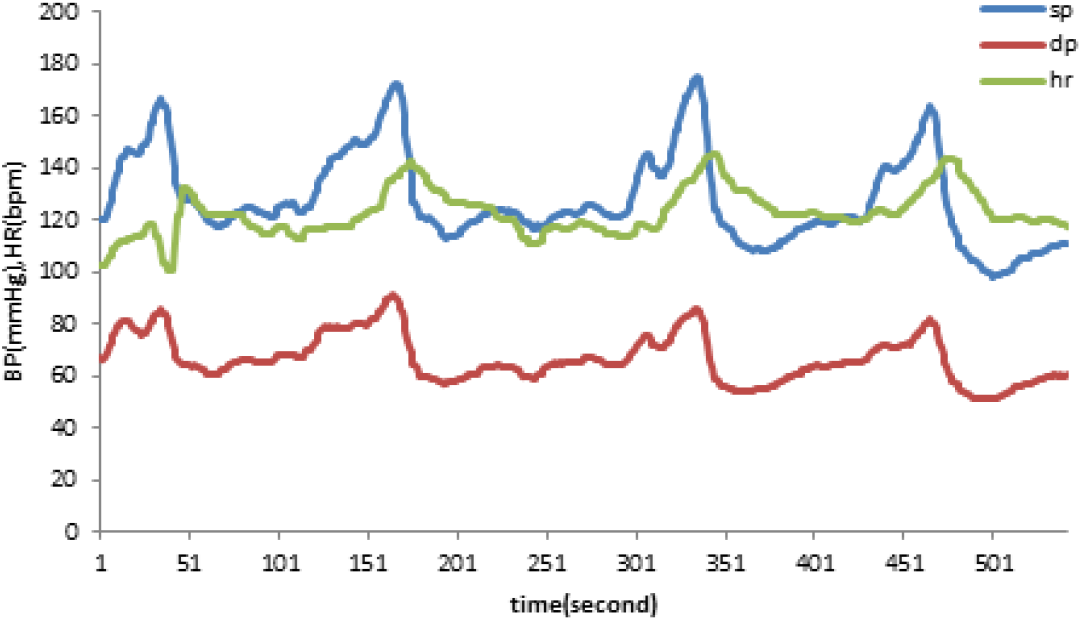
During intermittent exercise, BP and HR show continuous dynamic changes. During convalescence, SBP gradually decreased significantly, while HR did not show a corresponding obvious change pattern.

Kinetic analysis of SBP recovery curve showed that the change trend of SBP recovery curve was highly consistent with the double exponential decay model (R2 > 0.89), which accorded with the biphasic physiological mechanism mediated by neuroregulation and humoral regulation. The results of this study provide new insights for further understanding the recovery model of cardiovascular system in well-trained athletes after exercise.

In the case of constant resistance, no statistically significant difference in HR was observed between different treading frequencies (e.g. 40 rpm and 100 rpm). As shown in Fig. 2 (a), exercise duration is the main factor affecting HR changes. After HIE, the HR will continue to rise. Fig. 2 (b) shows the reaction of a weak subject: the HR drops rapidly after short-term exercise, while the HR still rises further after stopping exercise after long-term exercise, and the recovery period is significantly prolonged.

**Fig. 2.**
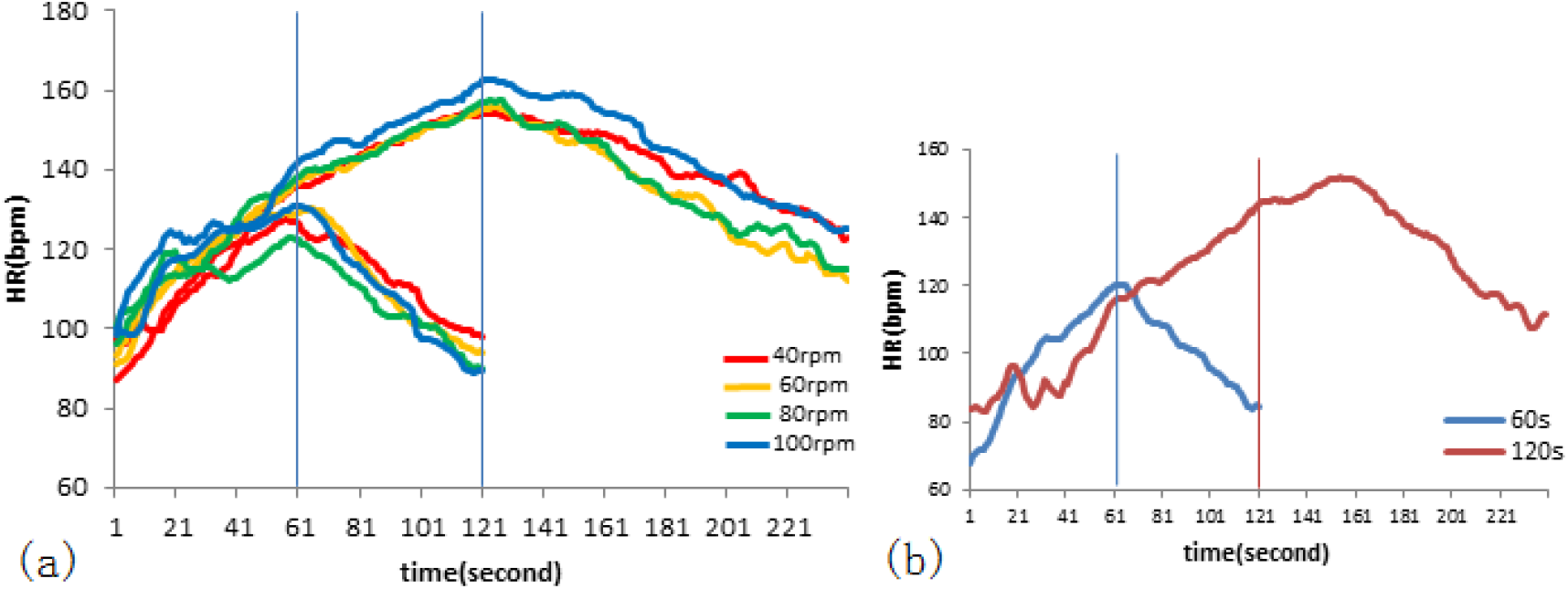
HR response under different load conditions. (a) The difference of HR between 60 seconds and 120 seconds of pedaling at different frequencies under 150W load. Under moderate load, the HR drops immediately after exercise, and there is a positive correlation between HR and exercise load, and the duration of exercise determines the maximum HR level. (b) The difference of HR between 60 seconds and 120 seconds of pedaling at different frequencies under 150W load. After the relatively HIE, the HR of some subjects continued to rise for a period of time after the exercise stopped.

After strenuous exercise, the HR continues to rise, while the BP drops rapidly. As shown in Fig. 3 (a), when the HR reaches its peak and the BP also drops to its valley, HR begins to drop, while SBP rises again. Between point B (movement stop) and point D (HR and BP trajectories re-meet), the decrease of SBP and the increase of HR show a highly consistent corresponding relationship, which indicates that there is physiological coupling between them. This close interaction suggests that the body can maintain hemodynamic stability through a compensatory mechanism during the recovery period after exercise.

**Fig. 3.**
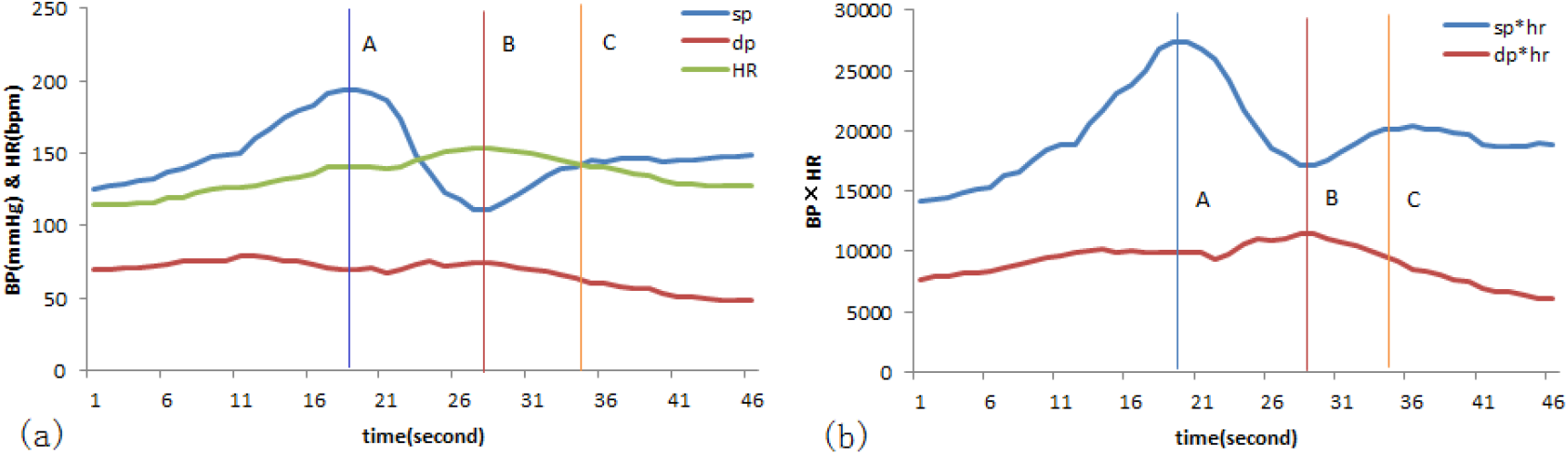
The BP-HR changes of a subject during a 17-second pedaling session, where line A indicates the end of pedaling, line B marks the time when BP drops to its lowest point and HR rises to its highest point, and line C indicates the time when BP rises again and HR intersects. (a) shows the segmented inflection points of each parameter change, and (b) presents the segmented calculation of EE rate and amount based on the segmentation in (a).

In the tests of running 60 meters and 100 meters on the athletics track, it is observed that: for the same participant, the HRmax during the 100-meter run is higher than that during the 60-meter run; after the 100-meter run, the increase in HR is greater compared to after the 60-meter run, and the duration of continued HR increase is longer after the 100-meter run than after the 60-meter run; during the 60-meter run, the HR recording line is steeper than during the 100-meter run, indicating that the derivative value of the fitting function of heart rate changes during the 60-meter run is larger. It can be inferred that for the participants in the test, the power output during the 60-meter run is higher, and more physical energy is consumed per unit time; during the 100-meter run, a greater amount of work is done.

Based on the PEE measured by DLW method, we obtained that the average PEE without moderate or HIE within a day is approximately 2030 Kcal. According to the differences in BP and HR between night and day, the PEE during 12 hours of the daytime is about 1050 Kcal, which calculates to an average PEE of 1.46 Kcal every minutes. It is correlated with the integral value of the BP-HR product fitting function in the corresponding time period, so as to obtain the PEE in this time period expressed by integral.

## 5 Discussion

Regardless of the form of exercise, for a specific individual, as long as the exercise intensity exceeds a certain value, the HR will continue to rise for a period of time after exercise ends, while BP will drop rapidly. After several repetitions of rising and falling with gradually decreasing amplitude, BP and HR tend to stabilize. This reflects the mutual coordination between BP and HR to maintain the body’s basic needs, and also provides a more solid theoretical basis for judging PEE based on BP-HR, putting forward new requirements for monitoring exercise-induced overtraining and ensuring exercise safety.

### 5.1 Evaluation method of high-intensity PEE

From the physiological mechanism of PEE detected by BP + HR, it is known that the coupling between hemodynamics and fluctuations in PEE, although HR decouples from metabolic rate during HIE, when combined with BP, it is not limited by HIE, reflecting consistency with metabolism. The quantitative representation of PEE during this period can be obtained through the integral change of BP × HR, and its mathematical expression is:

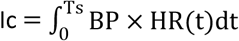

Where Ts represents the total duration for calculating the BP × HR integral.

Due to fluctuations in BP, the mean arterial pressure is generally calculated by adding one-third of the pulse pressure to the DBP. The mean arterial pressure can serve as a basis for calculating PEE during rest. However, during exercise, the HR increases, affecting the duration of systole and diastole, particularly impacting diastole more significantly, leading to changes in the ratio between systole and diastole, and consequently altering the mean arterial pressure. This change is quite complex; for the sake of discussion, we assume the mean arterial pressure to be constant.

Therefore, in Fig. 4 (a), the change in mean arterial pressure is approximately represented by line B, which is roughly parallel to DBP × HR. During the recovery period after exercise, the BP and HR multiplier exhibit multiple amplitude-decreasing fluctuations. We first calculate the integral from the start of exercise to the moment when the BP and HR multiplier first reach their lowest point. In Fig. 4 (a), on both sides of line A represents the process of exercise and the recovery process respectively. The high EE during the recovery process is caused by exercise and should be counted as part of the exercise EE. Therefore, the area above Line B and between SBP × HR represents the EE caused by exercise.

**Fig. 4.**
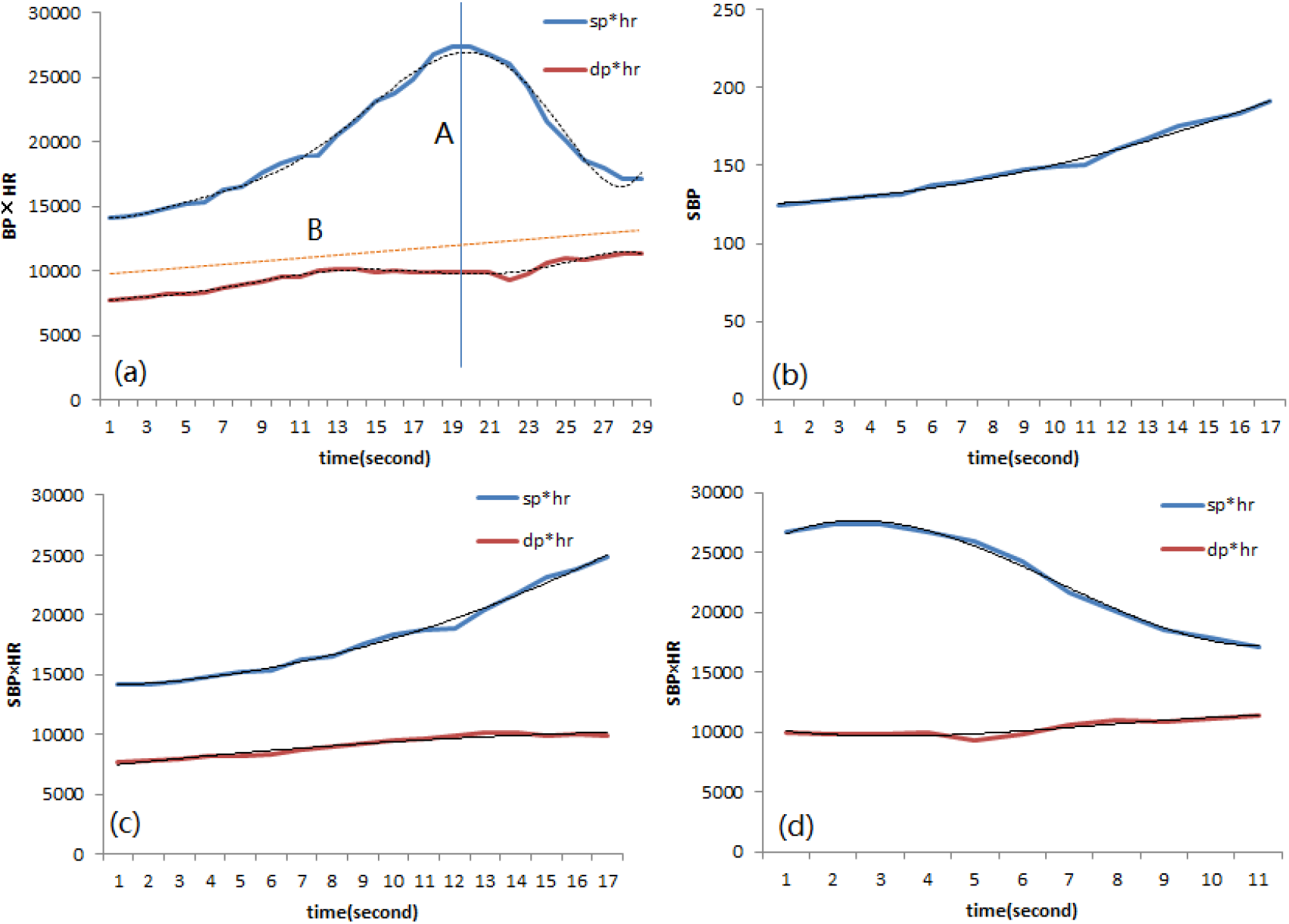
Calculation of EE and rate, where (a) the fitted function of BP×HR changes during the subject’s 17 seconds of pedaling and 12 seconds of rest, A line indicates the moment when BP×HR reaches its maximum, and F line represents the systolic BP level during the resting period; (b) the fitted function of SBP changes during the subject’s 17 seconds of pedaling; (c) the fitted function of BP×HR changes during the subject’s 17 seconds of pedaling; (d) the fitted function of BP×HR changes when the subject’s BP reaches its lowest point after pedaling.

In order to reduce the computational complexity, the motion process and the recovery process can be calculated by function integration respectively, as shown in Fig. 4 (c) and (d), based on the monotone increasing and monotone decreasing of continuous functions. The rate of PEE in a short period of time can be obtained by dividing the total amount of PEE in this period of time by the length of time. If we need to obtain the dynamic PEE rate, in Fig. 4(b), we can solve the differential equation constructed by the SBP fitting function in this period.

#### 5.1.1 The overall calculation of PEE during the exercise process

To achieve a higher goodness of fit, the integral of the fitting function for SBP × HR in Fig. 4(a) was performed using a high-order polynomial, obtaining the EE during this motion process. Before comparing it with the EE values measured by the DLW method, there were only quantitative standards, not specific energy expenditure values.

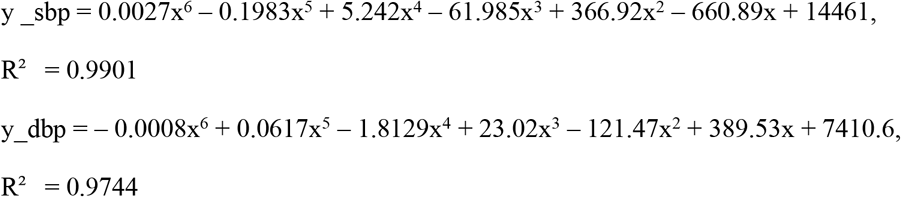

The product of mean arterial pressure and heart rate represents cardiac output during the resting period, which is the basal metabolism(bm).

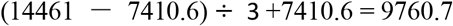

The difference between SBP and mean arterial pressure is caused by exercise.

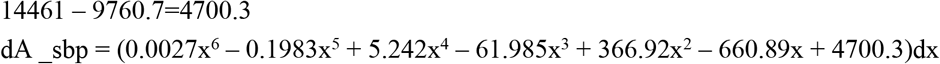

Integrals in T ∈ [0, 29]:

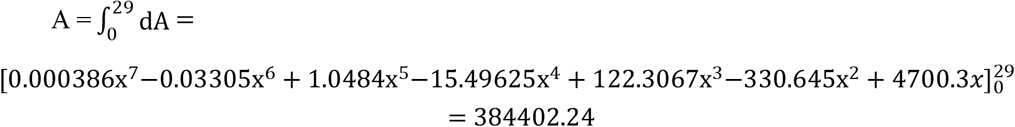

In Fig. 4(a), the area to the left of line A, above line B, and between SBP×HR represents the EE1 during exercise, while the area to the right of line A, above line B, and between SBP×HR represents the EE2 after exercise. The sum of EE1 and EE2 is the TEE caused by exercise. Combining these two parts for calculation requires a polynomial determined by goodness of fit, which involves significant computational effort. Therefore, calculating the exercise process and recovery process separately can reduce computational intensity.

On both sides of line A, the changes in SBP×HR are fitted with quadratic or cubic polynomials, which meets the requirements, in Fig.4(c) and Fig.4(d).

#### 5.1.2 The process of movement

In fact, the pedaling time was 17 seconds, but the maximum value of SBP×HR occurred at the 19th second. Therefore, when calculating SBP × HR, it is calculated based on 19 seconds, but when calculating the ratio of work time to recovery time, it is based on 17 seconds. The integral A of the difference between the SBP fitting function and the mean arterial pressure fitting function over the interval T ∈ [0,19] equals EE1, in Fig. 4(c).

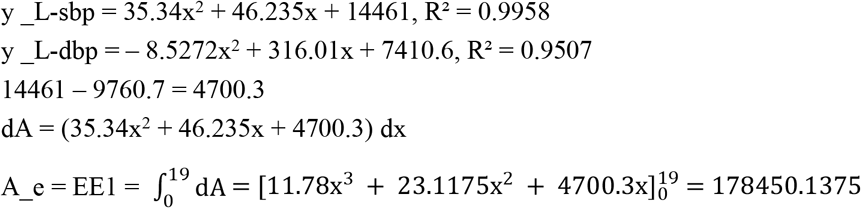

#### 5.1.3 The recovery process (until the end of the first wave’s fluctuation)

The integral A of the difference between the SBP fitting function and the mean arterial pressure fitting function over the interval T ∈ [0,10] equals EE2,in Fig. 4(d).

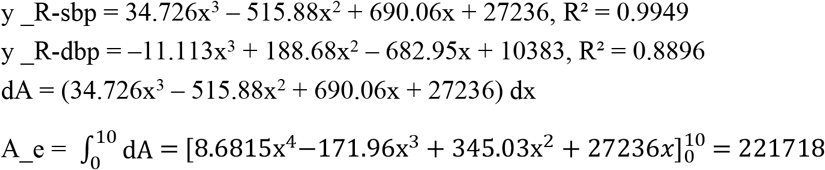

The area between the B line and the DBP fitting function represents the EE during the resting phase.

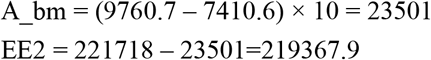

#### 5.1.4 The meaning of calculating numerical values

Due to the gap between segmented simulation and overall simulation of the function, the results of the two algorithms are not exactly the same, but they can illustrate the nature of the problem.

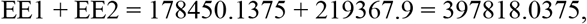

this value is approximately equal to 384402.

In addition, from (9760.7–7410.6) × 17 = 39951.7,the basal metabolic rate during a 17-second exercise period is approximately equal to 39951.7,therefore, the EE caused by physical activity is approximately 10 times that of basal metabolism. From the proportion of EE due to movement, one can roughly infer the participent’s level of tension and physical ability.

From the PEE of 1.46 Kcal during a one-minute quiet period, it can be derived that the PEE during one minute of HIE is 14.6 Kcal. Therefore, during the 17-second HIE period, PEE was 4.137 kilocalories, equivalent to 397818 or 384402 SBP×HR product integral units, that is, 93-96 units are equivalent to 1 calorie of energy.

#### 5.1.5 Calculation of short-term EE rate

The short-term EE rate determines the power of human work, as shown in Figure 4(b), obtained from the differential equation of the SBP fitting function.

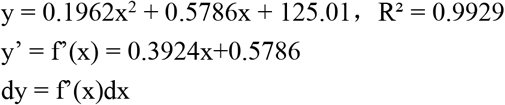

Since the exercise time is 17 seconds, therefore,

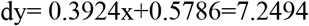

Through the value of the differential equation, a person can be judged his power output. However, there are currently no research results on sub-minute level PEE, so the PEE rate within a few seconds cannot be compared.

### 5.2 Rationality of the evaluation method

The TEE of three days and EE of each day were measured by DLW technique. Based on the TEE and EE values, it was found that there was a statistically significant correlation between SBP × HR and PEE in the same period (P < 0.01). Furthermore, the time integral ratio of SBP × HR realizes the segmented decomposition of TEE based on physiological process. The results not only have statistical stability, but also reflect the internal physiological coupling mechanism between cardiovascular response and metabolic demand, which shows significant biological rationality.

The findings of this study show that the time integral of BP × HR provides an accurate and highly practical measure for estimating EE during HIE. This index shows good robustness and adaptability in multi-mode sports scenes, and effectively overcomes the limitations of traditional univariate model under extreme physiological load.

In addition, the dynamic changes of SBP were highly consistent with the exercise intensity (R > 0.85)[3]. Specifically, during HIE, the EE rate can be reliably predicted according to the change rate of SBP, and the error range is controlled within 8%. This discovery provides a new physiological basis for real-time monitoring of exercise intensity, and has important value in high-demand application scenarios such as competitive sports and military training.

This study shows that the energy estimation method based on multivariate cardiovascular parameter integration is rigorous in methodology and superior to traditional models, which lays a solid theoretical and empirical foundation for the development of the next generation wearable monitoring system. Future research can focus on expanding sample diversity, optimizing real-time signal processing algorithms, and promoting the ecological application of this technology in real training environment.

## 6 Limitations

This study has limited participants to young adults around 19 years old and is related to sports training, which results in insufficient data representativeness. Additionally, the operational difficulties of invasive BP monitoring make it impractical for large-scale data collection; non-invasive BP monitoring has a low frequency, leading to the omission of many characteristic data points for sub-second level BP changes.

## 7 Practical implications

There is content in a textbook of exercise physiology related to calculating PEE, which introduces measuring PEE during 5 minutes of pedaling at a constant power using indirect calorimetry[93]. Our conclusion is similar to this, both obtaining the result that exercise PEE is approximately 10 times that of resting PEE. However, pedaling for 5 minutes cannot be maximum exercise intensity, so it still cannot be used as a basis for analyzing PEE during HIE lasting several seconds.

The credibility of the data results of this study is limited due to the presence of function fitting and systematic errors. However, with corrections made using large sample data, the reliability of the results is bound to improve. The purpose of this study is to construct a personalized energy expenditure model rather than pursuing uniform data for the group, thus the validity and credibility of the research results will be guaranteed. Thanks to the results of this study, high-frequency non-invasive BP measurement has become the focus of current equipment development, and in the future, the quantification of physical energy consumption will no longer avoid the need for high-intensity exercise.

## 8 Conclusions

Under the current research background of exercise physiology, it is still technically challenging to accurately quantify energy expenditure (EE) during short-term high-intensity exercise (HIE). In order to solve this problem, DLW method is used as the gold standard for measuring EE. By measuring the total energy expenditure (TEE) in three consecutive days, the comprehensive metabolic data including various exercise intensities (including high-intensity intermittent activities) were obtained. During this period, high-frequency invasive BP and HR monitoring was carried out simultaneously, and the time integral value of SBP and HR product (SBP × HR) was calculated systematically. Comparing the measured TEE and EE values with those measured by the multiplier integral method of BP and HR in the same time period, it was found that the ratios of them were highly consistent (r > 0.89), which indicated that the comprehensive cardiovascular parameters had high substitution with TEE measured by DLW, and proved that they had credible physiological significance in evaluating total energy output.

This study further proposes that the SBP × HR integral value corresponding to a specific HIE can be effectively used to estimate the EE of the exercise; The EE per unit time in a certain period of time is the rate of EE in that period, which can be calculated by dividing the TEE by time, and can also be obtained by solving the differential equation based on the SBP fitting function in that period.

This method achieves reasonable traceability from TEE to segment EE and proposes real-time monitoring of cardiovascular parameters during HIE to assess EE, as well as high-frequency non-invasive dynamic BP monitoring, providing new ideas and methodological support for applications in related fields.

## References

1. B. Ainsworth. E., et al. (2011). 2011 Compendium of Physical Activities: a second update of codes and MET values. Medicine & Science in Sports & Exercise, 43 (8), 1575–1581. DOI: 10.1249/MSS.0B013E31821ECE12

2. C. Garber. E., et al. (2011). Quantity and quality of exercise for development and maintaining cardiorespiratory, musculoskeletal, and neuromotor fitness in parallel healthy adults.Guidance for prescribing exercise. Medicine & Science in Sports & Exercise, 43 (7), 1334–1359. DOI: 10.1249/MSS.0B013E318213FEFB

3. Tanaka, H., Monahan, K. D., & Seals, D. R. (2001). Age-predicted maximum heart rate revised. Journal of the American College of Cardiology, 37 (1), 153–156. DOI: 10.1016/S0735-1097(00)01054-8

4. Borg, G. A. (1982). Psychophysical bases of perceived extinction. Medicine & Science in Sports & Exercise, 14 (5), 377–381.

5. Laursen, P. B., & Jenkins, D. G. (2002). The scientific basis for high-intensity interval training: optimizing training programs and maximizing performance in highly trained endurance athletes. Sports Medicine, 32 (1), 53–73.

6. Søgaard, K., et al. (2017). High-intensity exercise for workplace health promotion: a systematic review and meta-analysis. Scandinavian Journal of Work, Environment & Health, 43 (4), 301–311.

7. M. Gibala. J., et al. (2012). Physiological adaptations to low-volume, high-intensity interval training in health and disease. The Journal of Physiology, 590 (5), 1077–1084. DOI: 10.1113/jphysiol.2011.224725

8. MacInnis, M. J., & Gibala, M. J. (2017). Physiological adaptations to interval training and the role of excise intention. The Journal of Physiology, 595 (9), 2915–2930. DOI: 10.1113/JP273196

9. Wisløff, U., et al. (2007). Superior cardiovascular effect of interval training versus modern excise in heart failure patients: a randomized controlled trial. Circle, 115 (24), 3086–3094. DOI: 10.116/circulationaha.106.675041

10. Buchheit, M. & Laursen, P. B. (2013). High-intensity interval training, solutions to the programming puzzle. Part I: cardiopulatory emphasis. Sports Medicine, 43 (5), 313–338. DOI: 10.1007/s40279-013-0029-x

11. Aagaard, P., et al. (2003). Increased rate of force development and neural drive of human skeletal music following persistence training. Journal of Applied Physiology, 95 (6), 2318–2326.

12. G. Millet. P., et al. (2011). Neurophysiological adaptations to endurance and interval training. Medicine & Science in Sports & Exercise, 43 (5), 313–338.

13. Girard, O., et al. (2011). Neuromuscular fatigue in racquet sports. Neuromuscular Aspects of Sport Performance, 1–28.

14. J. Lauersen. B., et al. (2018). The effectiveness of excise intervals to advance sports injuries: a systematic review and meta-analysis of random controlled trials. British Journal of Sports Medicine, 48 (11), 871–877. DOI: 10.1136/bjsports-2013-092538

15. W. Meeuwisse. H., et al. (2007). A dynamic model of etiology in sport injection: the recursive nature of risk and cause. Clinical Journal of Sport Medicine, 17 (3), 215–219.

16. Eriksson, H., & Lindgren, G. (2014). Physical work in mining and mining construction. Arbete och Ha ® lsa, 48 (4), 1–48.

17. Bao, S., & Silverstein, B. (2005). Estimation of metallic energy expansion in manual material handling jobs. Journal of Electromyography and Kinesiology, 15 (1), 1–9.

18. Torp, S., et al. (2013). The impact of mandatory physical activity on cardiovascular disease and all-cause morality: a systematic review. Scandinavian Journal of Work, Environment & Health, 39 (4), 327–338.

19. W. Grooten. J. A., & Wåhlin, C. (2006). J. International Journal of Industrial Ergonomics, 36 (1), 1–12.

20. B. Nindl. C., et al. (2017). Physiological, psychological, and performance correlations of military operational stress. Medicine & Science in Sports & Exercise, 49 (5S), 1–12.

21. Henning, P. C., et al. (2011). Physiological and psychological responses to a high-intensity interval training program designed for special operations forces. Journal of Strength and Conditioning Research, 25 (3), 668–677.

22. K. Burgomaster. A., et al. (2008). Similar metabolic adaptations during exercise after low volume sprint interval and traditional endurance training in humans. The Journal of Physiology, 586 (1), 151–160.

23. MacInnis, M. J., & Gibala, M. J. (2017). Physiological adaptations to interval training and the role of excise intention. The Journal of Physiology, 595 (9), 2915–2930. DOI: 10.1113/JP273196

24. N. Vollaard (1800). B. J., et al. (2017). Systematic review and meta-analysis of the effect of high-intensity interval training on maximum oxygen uptake and mitochondrial function. Medicine & Science in Sports & Exercise, 49 (6), 1057–1065.

25. LaForgia, J., Withers, R. T., & Gore, C. J. (2006). Effects of excise intensity and duration on the exception post-excise oxygen consumption. Journal of Sports Sciences, 24 (12), 1247–1264.

26. A. Knab. M., et al. (2011). A 45-minute vigorous exercise bout increments metabolic rate for 14 hours. Medicine & Science in Sports & Exercise, 43 (9), 1643–1648. DOI: 10.1249/MSS.0B013E3182118891

27. Børsheim, E., & Bahr, R. (2003). Effect of excise intention, duration and mode on post-excise oxygen consumption. Sports Medicine, 33 (14), 1037–1060.

28. Sigal, R. J., et al. (2014). Effects of aerobic training, persistence training, or both on glycemic control in type 2 diabetes: a randomized trial. Annals of Internal Medicine, 161 (2), 121–130.

29. E. Richter. A., & Hargreaves, M. (2013). Exercise, GLUT4, and Skeletal Muscle Glucose Uptake. Physiological Reviews, 93 (3), 993–1017. DOI: 10.1152/physrev.00038.2012

30. Jelleyman, C., et al. (2015). The effects of high-intensity interval training on glucose regulation and insulin resistance: a meta-analysis. Obesity Reviews, 16 (11), 942–961. DOI: 10.1111/obr.12317

31. Colberg, S. R., et al. (2016). Physical Activity/Exercise and Diabetes: A position statement of the American Diabetes Association. Diabetes Care, 39 (11), 2065–2079. DOI: 10.2337/DC16-1728

32. B. Beck. R., et al. (2017). Exercise and Sports Science Australia (ESSA) position statement on excise prescription for the prevention and management of osteoporosis. Journal of Science and Medicine in Sport, 20 (5), 438–445. DOI: 10.1016/j.jsams.2016.02.001

33. J. Ramos. S., et al. (2015). The impact of high-intensity interval training versus modern-intensity continuous training on variable function: a systematic review and meta-analysis. Sports Medicine, 45 (5), 679–692. DOI: 10.1007/s40279-015-0321-z

34. Wiss øff, U., et al. (2007). Superior cardiovascular effect of interval training versus modern excise in heart failure patients: a randomized controlled trial. Circle, 115 (24), 3086–3094.

35. Eijsvogels, T. M. H., Fernandez, A. B., & Thompson, P. D. (2016). Are there deleterious cardiac effects of acute and chronic endurance exercise? Physiological Reviews, 96 (1), 99–125.

36. La Gerche, A., & Heidbuchel, H. (2014). Can intentional exercise harm the heart? Insights from a study of elite athletes. Circle, 130 (12), 987–991.

37. Bahr, R., & Krosshaug, T. (2005). Understanding injection mechanisms: A key component of adventing injuries in sport. British Journal of Sports Medicine, 39 (6), 324–329.

38. Powers, S. K., & Jackson, M. J. (2008). Exercise-induced oxidative stress: cellular mechanisms and impact on muscle force production. Physiological Reviews, 88 (4), 1243–1276. DOI: 10.1152/physrev.00031.2007

39. Nieman, D. C., & Wentz, L. M. (2019). The compelling link between physical activity and the body’s defense system. Journal of Sport and Health Science, 8 (3), 201–217. DOI: 10.1016/j.jshs.2018.09.009

40. D. Tietze. C., & Borchers, J. (2014). Exertional rhabdomyolysis in the athletic: a clinical review. Sports Health, 6 (4), 336–339. DOI: 10.1177/1941738114523544

41. Nieman, D. C., & Wentz, L. M. (2019). The compelling link between physical activity and the body’s defense system. Journal of Sport and Health Science, 8 (3), 201–217. DOI: 10.1016/j.jshs.2018.09.009

42. Thompson, P. D., et al. (2007). Exercise and Acute Cardiovascular Events: Placing the risks into perspective. Medicine & Science in Sports & Exercise, 39 (5), 886–897.

43. K. Harmon. G., et al. (2015). Pathogeneses of Sudden Cardiac death in competitive athletes. Circle: Arrhythmia and Electrophysiology, 8 (1), 218–229. DOI: 10.1161/CIRCEP.113.001376

44. Bahr, R., & Holme, I. (2003). Risk factors for sports injuries-a methodological approach. British Journal of Sports Medicine, 37 (5), 384–392.

45. Junge, A., & Engebretsen, L. (2009). Injury survey in multi-sport events: the International Olympic Committee approach. British Journal of Sports Medicine, 43 (6), 413–421. DOI: 10.1136/bjsm.2008.046631

46. D. Schoeller. A. (1988). Measurement of energy expansion in free-living humans by using doubly labeled water. Journal of Nutrition, 118 (11), 1278–1289.

47. K. Westerterp. R. (2017). Doubly labeled water assessment of energy expansion: principle, practice, and promise. European Journal of Applied Physiology, 117 (7), 1277–1285.

48. IAEA (International Atomic Energy Agency). (2009). Assessment of Body Composition and Total Energy Expansion in Humans Using Stable Isotope Technologies. IAEA Human Health Series No. 3.

49. Pontzer, H., et al. (2016). Constrained Total Energy Expansion and Metabolic Adaptation to Physical Activity in Adult Humans. Current Biology, 26 (3), 410–417. DOI: 10.1016/j.cub.2015.12.046

50. Pontzer, H., et al. (2021). Daily energy expansion through the human life course. Science, 373 (6556), 808–812. DOI: 10.1126/SCIENCE.ABE5017

51. Trabulsi, J., & Schoeller, D. A. (2001). Evaluation of the double labeled water method against the double labeled water method in obese subjects. American Journal of Physiology-Endocrinology and Metabolism, 281 (2), E357–E367.

52. D. Schoeller. A., & Hnilicka, J. M. (1996). Reliability of the double labeled water method for the measurement of total daily energy expansion in free-living subjects. Journal of Nutrition, 126 (1), 348S–354S.

53. J. Speakman. R. (1998). The history and theory of the double labeled water technique. American Journal of Clinical Nutrition, 68 (4), 932S–938S.

54. Yao, M., et al. (2002). Energy requirement and basic metal rate of Chinese young men. Asia Pacific Journal of Clinical Nutrition, (7), S226–S230

55. Yosuke, F., et al. (2023). Validation of body fat measurement using dual-energy X-ray absorptiometry and bioelectrical impedance analysis by dual-label water method in Japanese older adults. Geriatrics & Gerontology International, 23 (5), 381-387. DOI: org/10.1111/ggi.14568

56. Elia, M., et al. (1992). The measurement of carbon dioxide production using 13C bicarbonate in the newborn. Pediatric Research, 31 (3), 255–260.

57. Schadewaldt, P., et al. (1997). Application of isotopic-selective nonspecific infrared spectrometry (IRIS) for evaluation of 13C bicarbonate kinetics in humans. Clinical Chemistry, 43 (3), 480–486.

58. Van der Schoor, S. R., et al. (2007). The use of the 13C bicarbonate technique for the evaluation of energy metabolism in infants: a review. Journal of Pediatric Gastroenterology and Nutrition, 44 (2), 142–152.

59. Ekelund, U., et al. (2001). Physical activity assisted by activity monitor and doubly labeled water in children. Medicine & Science in Sports & Exercise, 33 (2), 275–281.

60. Ekelund, U., et al. (2006). Criterion-related validity of the last 7-day, short form of the International Physical Activity Questionnaire in Swedish adults. Public Health Nutrition, 9 (2), 258–265. DOI: 10.1079/PHN2005840

61. Brage, S., et al. (2015). Comparison of PAEE from combined and separate heart rate and movement models in children. Medicine & Science in Sports & Exercise, 47 (6), 1311–1319.

62. Brage, S., & Westgate, K. (2012). How to measure physical activity in real life? Obesity Reviews, 13 (2), 1–3.

63. S. Intille. S., et al. (2012). New developments in sensing and context-awareness for health and fitness. IEEE Pervasive Computing, 11 (4), 24–31.

64. J. Steeves. A., et al. (2015). Pattern recognition methods for estimating physical activity using body-word sensors. IEEE Journal of Biomedical and Health Informatics, 19 (4), 1389–1396.

65. W. Welch. A., et al. (2017). Accuracy of a combined heart rate and accelerometer sensor for measuring energy expansion in free-living adults. Journal of Science and Medicine in Sport, 20 (7), 675–680.

66. Ekelund U., Yngve A., Westerterp K., et al. Energy Expenditure Assessed by Heart Rate and Doubly Labeled Water in Young Athletes. Medicine and Science in Sports and Exercise, 2002, 34 (8): 1360–6.

67. Feng J. A New Technology for Measuring Energy Expenditure: The Doubly Labeled Water Method. Chinese Journal of Sports Medicine, 1996, (03): 203–205.

68. D. Macfarlane. J. (2001). Automated metabolic gas analysis systems: a review. Sports Medicine, 31 (12), 841–861.

69. D Bishop. J., et al. (2014). The limits of the constant load test for the assessment of cycling performance. European Journal of Applied Physiology, 114 (2), 345–362.

70. Achten, J., & Jeukendrup, A. E. (2003). Heart rate monitoring: applications and limits. Sports Medicine, 33 (7), 517–538.

71. S. Crouter. E., et al. (2018). Validity of the ActiHeart for the Assessment of Energy Expenditure in Adults. Medicine & Science in Sports & Exercise, 50 (6), 1335–1342.

72. Benedetto, S., et al. (2018). A review of wearable and non-wearable technologies for monitoring human physiological signals. Sensors, 18 (9), 3,084.

73. An, H. S., & Jones, G. C. (2018). A Review of Wearable Motion Tracking Systems for Sports and Rehabilitation. Journal of Biomedical Engineering and Medical Imaging, 5 (2), 1–10.

74. Brage, S., et al. (2004). Hierarchy of individual calibration levels for heart rate and accelerometry to measure physical activity. Journal of Applied Physiology, 97 (2), 697–704.

75. Winkert, K., et al. (2021). The need for an differentiated approach to energy expansion period in athletes. European Journal of Sport Science, 21 (5), 709–719.

76. Gonzalez-Alonso, J., et al. (2008). Cardiovascular and metabolic responses to excise in the heat: a review. International Journal of Sports Medicine, 29 (8), 603–612.

77. J. Cuddy. S., et al. (2014). The effects of heat stress on the physiological responses to excise: a review. Journal of Strength and Conditioning Research, 28 (10), 2980–2995.

78. A. Guyton. C., & Hall, J. E. (2006). Textbook of medical physiology (11th ed.). Elsevier Saunders.

79. L. Rowell. B. (1993). Human cardiovascular control. Oxford University Press.

80. Chapman, C. L., De Castro, J. M., Ely, M. R., Johnson, B. D., & Minson, C. T. (2020). Blood pressure regulation VIII: Resistance vessel tone and implication for a prohypertensive effect of low-dose aspirin in women. American Journal of Physiology-Heart and Circular Physiology, 319 (1), H241–H255. DOI: 10.1007/s00421-013-2684-x

81. Joyner, M. J., & Casey, D. P. (2015). Regulation of incremented blood flow (hyperemia) to muscles during excise: a hierarchy of matching physiological needs. Physiological Reviews, 95 (2), 549–601.

82. M. Reeder. K., & Green, H. J. (2012). Muscle bloodflow during exercise: the limits of reductionism. Medicine and Science in Sports and Exercise, 44 (5), 813–817.

83. MS. Laughlin (1800’s). H., Davis, M. J., Secher, N. H., van Lieshout, J. J., Arce-Esquivel, A. A., Simmons, G. H.,… & Boushel, R. (2012). Peripheral circulation. Comprehensive Physiology, 2 (1), 321–447.

84. Saltin, B., & Mortensen, S. P. (2012). Inefficent functional sympathology is an overlooked cause of malperfusion in contracting skeletal music. The Journal of Physiology, 590 (24), 6269–6275. DOI: 10.1113/jphysiol.2012.241026

85. B. Wallin. G., & Sundloöf, G. (1982). Sympathy outflow to muscles during vasovagal syncope. Journal of the Autonomic Nervous System, 6 (3), 287–291.

86. Esler, M., et al. (1988). Information of human symmetrical nervous system activity from measures of norepinephrine turnover. Hypertension, 11 (1), 3–20.

87. Hofmann, P., & Tschakert, G. (2011). Intensity-and duration-based options to regulate endurance training. Frontiers in Physiology, 2, 71. DOI: 10.3389/fphys.2017.00337

88. Bernardi, E., et al. (2020). Heart Rate Variability in the Assessment of Exercise Intensity in Endurance Athletes: A Systematic Review. Annals of Biomedical Engineering, 49 (5), 1311–1326.

89. C. Vella. A., & Robergs, R. A. (2005). A review of the stroke volume response to priright excise in healthy subjects. British Journal of Sports Medicine, 39 (4), 190–195.

90. Brage, S., et al. (2007). Hierarchy of individual calibration levels for heart rate and accelerometry to measure physical activity. Journal of Applied Physiology, 103 (2), 682–692.

91. Höchsmann, C., et al. (2019). A Review of the Use of Combined Heart Rate and Movement Sensors to Estimate Energy Expenditure in Humans: The Emerging Role of Machine Learning. Sensors, 19 (19), 4137.

92. Garcia, S., et al. (2022). A Multimodal Sensing Platform for Interdisciplinary Training in Data-Rich Sports and Exercise Science Research. IEEE Sensors Journal, 22 (5), 4095–4107.

93. R. Wang, et al. Exercise Physiology. People’s Sports Press, 2007: 166–168. 978-7-5009-2309-1

